# Plasticity of threespine stickleback nuptial signals: laboratory manipulation of ambient light causes a shift in male reflectance patterns

**DOI:** 10.1101/2022.05.17.492337

**Authors:** Robert J. Scott

## Abstract

I caught courting male threespine stickleback (*Gasterosteus aculeatus*) in Hotel Lake, BC, Canada, and transferred them to one of three experimental lighting environments. Male signal reflectance was measured upon capture from the lake, at one-day post transfer to the experimental light, and at onset of courtship behaviour under artificial light. Male signals responded rapidly to the change in lighting environment following transfer from the wild to the artificial lighting environments and in different ways dependent upon the artificial light composition. Males under red-shifted light had higher signal reflectance in the short wavelength range of the spectrum and less reflectance in the medium and long wavelength range of the spectrum relative to the males in blue-green shifted and full-spectrum lighting. Given the relative prominence of the male stickleback nuptial signal in the behavioural ecology literature, the increasing evidence for the signal’s ambient light dependent expression, and the large body of literature based on experiments conducted under artificial light, it would be prudent to repeat some of the classic mate choice experiments under more realistic lighting conditions.

## Introduction

Coupling of signal and receiver properties is necessary for reliable information exchange in communication systems (Endler 1991, Hauser 1997). However, the efficacy of a signalling system is subject to variability in the signalling environment since signals must stand out relative their background and transmit well through the signalling medium (Endler 2000, van der Sluijs et al., 2011, Stevens 2013). Many species are comprised of populations that span a range of habitats that impact signal contrast and transmission (e.g. Ryan and Wilczynski 1991, Scott 2001, Fuller and Noa 2010, Barquerro et al. 2015, Giery and Layman 2017). The efficacy of a fixed signalling phenotype would vary among habitats, having higher efficacy in some conditions and reduced or lost efficacy in others (Via et al. 1995). Therefore, properties of communication systems should vary among habitats to maintain effectiveness (Endler 2000, van der Sluijs et al. 2011, Wiley 2013).

Visual signals in aquatic ecosystems are influenced by environmental conditions (Lythgoe 1979); light transmission and absorption are affected by material dissolved and suspended in the water, including the planktonic community (Wetzel 2001, Bracchini et al. 2005, V-Balogh et al. 2009, Szeto et al. 2011). Physical and ecological processes within lakes and their surrounding drainage basins influence the type and amount of light absorbing and scattering materials present in a lake (Pace and Cole 2002, Morris et al. 1995, Seekell et al. 2014). Variability among lake systems in the processes that influence dissolved and suspended material produces variability among lakes in optical properties. Variation of this sort has lead to population differences in mating signals across a number of fish species in diverse aquatic systems (*Poecilia reticulata*: Houde and Endler 1990; *Gasterosteus aculeatus*: Reimchen 1989, Scott 2001; *Gambusia bahamasminimus*: Giery and Layman 2017; *Lucania goodei*: Fuller and Noa 2010).

Male threespine stickleback produce a mosaic nuptial signal, comprised of a red opercular region (chin), bright blue iris and blue-green dorsum (McLennan and McPhail 1989, Rowland 1994). The red component of the signal is produced by the deposition of carotenoid pigments in the dermis (Wedekind et al. 1998, Pike et al. 2011; but see Reimchen 1981, Scott 2011) and variability among males in expression of this signal has been argued to indicate male carotenoid content and, since carotenoids are rare in nature, to reflect male quality (Barber et al. 2000, Milinski and Bakker 1990, Bakker and Mundwiler 1994, Kunzler and Bakker 2001, Pike et al. 2007a, b, 2011). However, aside from studies examining large changes in the nuptial signal (for example complete loss or evolution of melanism; Reimchen 1989; Scott 2001, 2011) there has been little attention to how the signal responds to environmental variability in lighting conditions.

The objective of the present study is to examine the response of the male threespine stickleback (*G. aculeatus*) nuptial signal following transfer from natural lighting conditions in the wild to three different artificial lighting conditions in the laboratory. Given the demonstrated importance of the nuptial signal in this species during courtship (e.g. Barber et al. 2000, Kunzler and Bakker 2000, Pike et al. 2007a, 2011) and the potential alteration of the efficacy of the signal under different lighting conditions (Reimchen 1989; Scott 2001), I predict that the signal will change between the natural and artificial settings and that the signal will diverge under each of the three artificial lighting conditions.

### The threespine stickleback system

The threespine stickleback is found in coastal marine, brackish, and freshwater systems of the Holarctic biogeographic realm (Bell and Foster 1994). This species is notoriously good at colonizing new habitats; most freshwater populations throughout the bulk of its distribution has arisen since the retreat of the most recent glaciation as a result of colonization by the long-present marine form (Bell et al. 2004, Lescak et al. 2015, Leaver and Reimchen 2012). Whereas morphological, physiological, life history and behavioural variability among marine and freshwater populations of this species have been examined and documented fairly extensively, population variability in the mating signal has by and large been ignored (but see Reimchen 1989, Scott and Foster 2000, Scott 2001, 2004, 2011, Candolin et al. 2016 for exceptions). Additionally, plasticity for morphological, physiological, and behavioural traits has been demonstrated in both ancestral (marine) and derived (freshwater) populations (for example morphology: Wund et al. 2008, McCairns and Bernatchez 2012, Leaver and Reimchen 2012; physiology: Gibbons et al. 2017, Morris et al. 2014; behaviour: Foster 1999, Foster 2013; Heuschele et al. 2012). Plasticity of the mating signal has received little attention (but see Hulslander 2003; Brock et al. 2017), This study examines impacts of varying ambient light expression of the red component of the signal. Stickleback mate choice experiments are most often conducted under artificial lighting (but see Scott 2004) and demonstrate the importance of the red component of the male signal in choice by females. However, if expression of the red signal is influenced by ambient light, then future stickleback mate choice studies should attempt to use natural lighting.

## Methods

### Experimental set-up

Nesting male stickleback in Hotel Lake, British Columbia, Canada (49°38’18” N, 124°02’45” W) were identified by a diver in the water and observed for several minutes to determine whether each was in the territory set-up, nest construction, courting, or parental phase of the reproductive interval. Males that were observed to actively court a gravid female were captured (using a dip net) and the reflectance spectra of each male’s opercular region was measured (Ocean Optics SD2000 spectrometer fitted with reflectance probe R-400 and calibrated with white standard WS-1) within 1.5 minutes of capture (Laurin and Scott 2011). Males were transported to the lab and held in 80 l aquaria and fed frozen brine shrimp *ad libidum* until commencement of the experiment (3 days). Males were randomly assigned to one of three lighting treatments and placed individually into 20 l aquariums. Each aquarium had washed sand on the bottom, an air stone, and natural nesting material (algae and detritus from the lake). I measured the reflectance spectrum of each male’s opercular region again after one day of acclimatizing to the new tank (Day 0). A gravid female was presented in a transparent container to each male daily for 30 minutes to initiate nesting behaviour. Each male was re-captured and I measured the spectral reflectance of his opercular region again once he had constructed a nest and was observed readily courting the gravid female. Males were euthanized (overdose of MS-222) following the final reflectance measurement.

### Lighting treatments

Tanks for each treatment were illuminated with 122 cm T8 32 Watt full spectrum fluorescent lamps (Maintenance Engineering model F32T8/VL8454/RS) suspended 40 cm above the tank. Lighting treatments were achieved by placing each fluorescent lamp in a clear polycarbonate tube (Roscosleeve model T8×48, Rosco Canada) containing a colour filter (Rosco Canada). Three lighting environments were produce: red shifted (Rosco Supergel #26 – Light Red), blue-green (Rosco Supergel #374 – Sea Green) and full-spectrum reduced intensity (Rosco Cinegel #3402 – Neutral Density).

The absolute irradiance of down-welling light at 47 nests in Hotel Lake and in each of the three lighting treatments was measured using a spectroradiometer (model SD2000, Ocean Optics, FL, USA) attached to a fibre optic probe fitted with a cosine corrector (model CC-3, Ocean Optics, FL, USA) held 10 cm above a male’s nest. Irradiance measurements in Hotel Lake were made within 1 hr on either side of solar noon. Irradiance distributions for Hotel Lake and each of the lighting treatments are presented in Figure 1.

**Figure 1.**
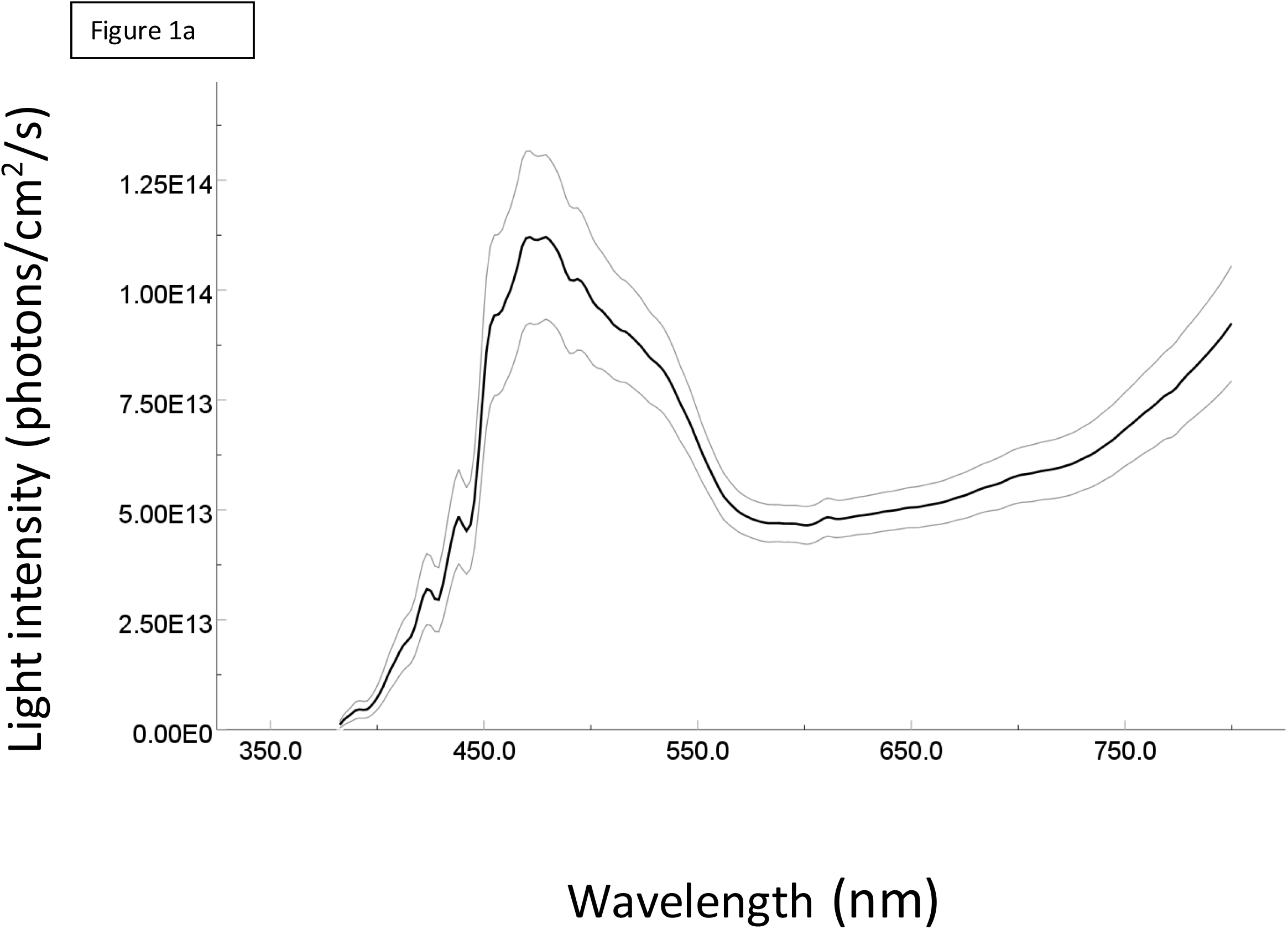
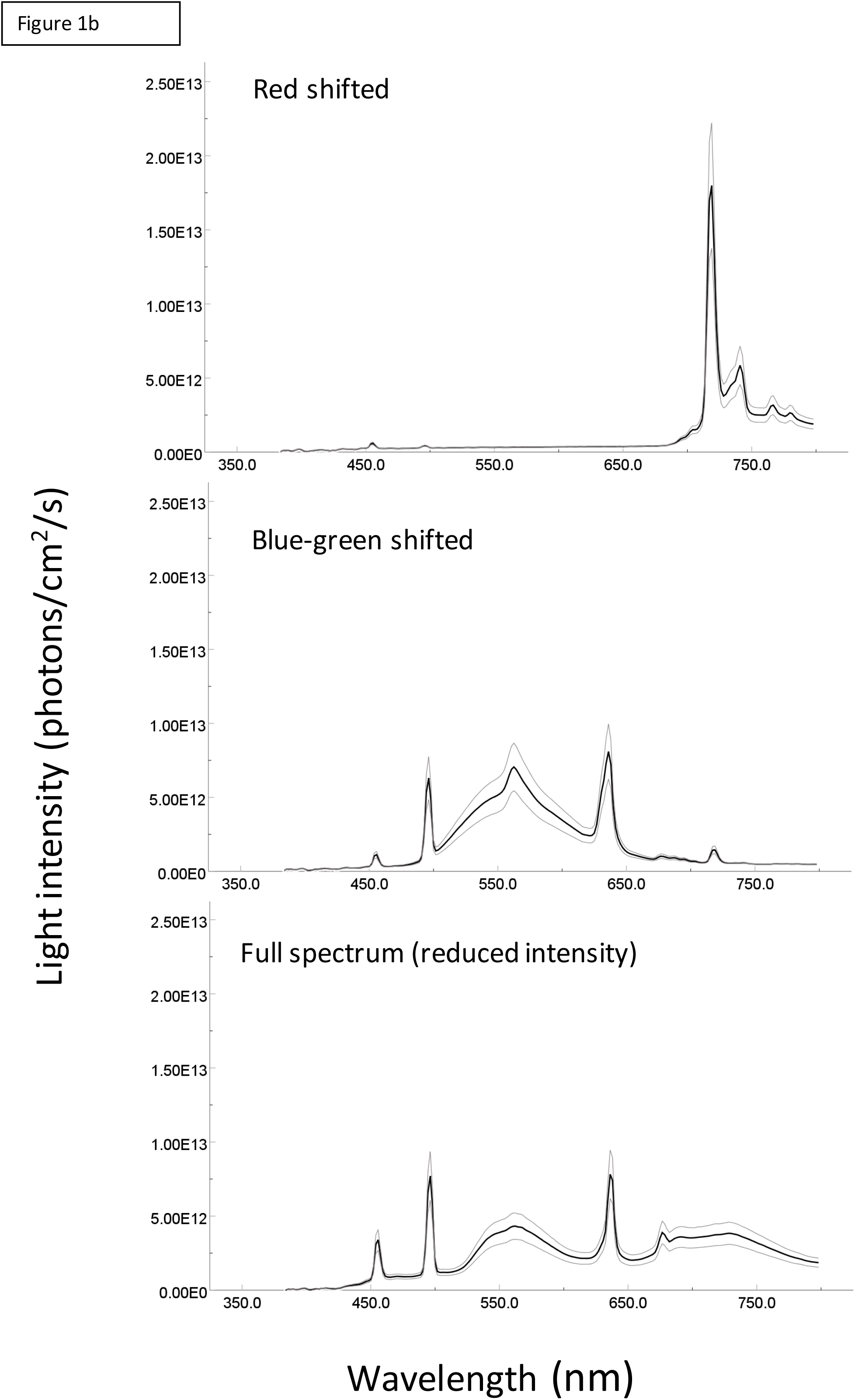
Spectral characteristics of (a) Hotel Lake and (b) each of the three lighting treatments. For each graph, the black line represents the mean and the gray lines represent the 95% confidence intervals.

### Characterizing male signal reflectance

Reflectance spectra were captured for wavelengths ranging from 339 – 1017 nm at 0.34 nm intervals, producing 2048 data points per individual. Subsequently, I reduced the number of data points to 208 by first truncating the range to 384 nm to 798 nm and averaging adjacent values at 2 nm intervals (Cuthill *et al*. 1999; Fuller 2002). Reduced data were then characterized by the relative intensity of short (384-490 nm), medium (492-580 nm), and long (582-798 nm) portions of the visible spectrum using the following equations (modified from Endler 1990);

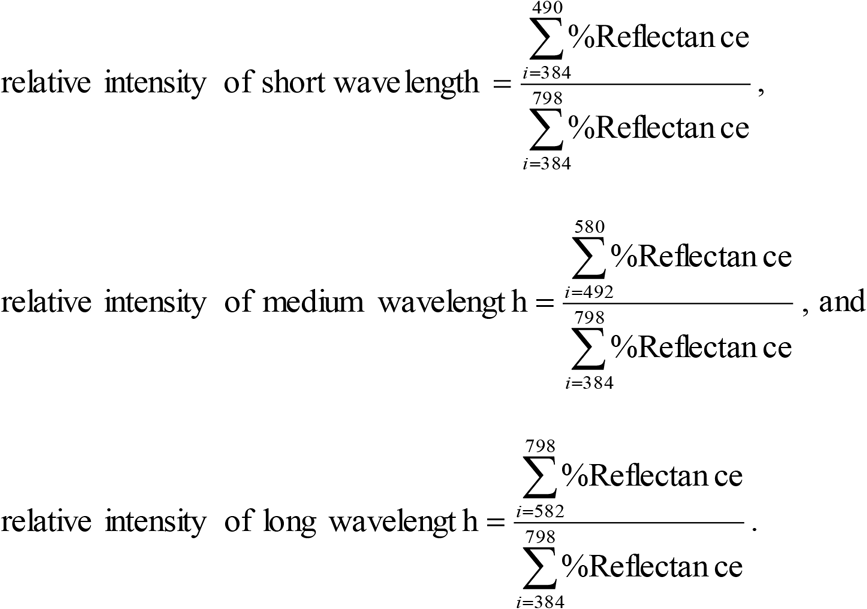

### Data analysis

I used mixed model ANOVA to examine the impacts of the three artificial lighting environments on signal expression since each individual was subjected to only one lighting treatment but measured repeatedly (at capture from the lake, at day 1 and again when actively courting a female) (Quinn and Keough 2004). I ran separate analyses for the short, medium and long wavelength components of the signal.

## Results

I captured and measured signals of 30 males that were observed courting females in the wild. All males constructed nests under the experimental conditions, but three males (two from the full spectrum and one from the blue-green shifted treatments) ceased nest construction at day four and five of the experiment. The remaining 27 individuals continued nest construction and began actively courting females between day 10 and 13 of the experiment.

Analyses revealed lack of sphericity for all three signal components (Mauchley’s W = 0.365, 0.696 and 0.715 for the short, medium and long wavelength components of the signal respectively; df = 2 and p < 0.05 for each). All subsequent analyses used the Greenhouse-Geisser corrected p-values (Quinn and Keough 2004). Signal reflectance in the short range of the spectrum was influenced by both the measurement day (Figure 2a; F_1.2, 29.4_ = 42.133, p < 0.005) and lighting treatment (F_2, 24_ = 7.706, p = 0.003). There may be an interaction between measurement day and lightning treatment (F_2.45, 29.35_ = 2.89, p = 0.062) with reflectance in the short range changing more in the full and blue-shifted treatments compared to the red-shifted treatment. Signal reflectance in the medium range of the spectrum was influenced by both the measurement day (Figure 2b; F_1.5, 36.823_ = 44.392, p < 0.005) and lighting treatment (F_2, 24_ = 5.316, p = 0.012). There is no interaction between these two variables (F_3.069, 36.823_ = 1.145, p = 0.345) suggesting that the changes to the signal caused by lighting environment are the same over time. Signal reflectance in the long range of the spectrum was influenced by both the measurement day (Figure 2c; F_1.557, 37.363_ = 17.730, p < 0.005) and lighting treatment (F_2, 24_ = 4.158, p = 0.028). Again, there may be an interaction between these two variable (F_3.114, 37.363_ = 2.172, p = 0.086) with the full spectrum and blue-shifted treatments producing greater change in long-wave reflectance compared to the red-shifted treatment.

**Figure 2.**
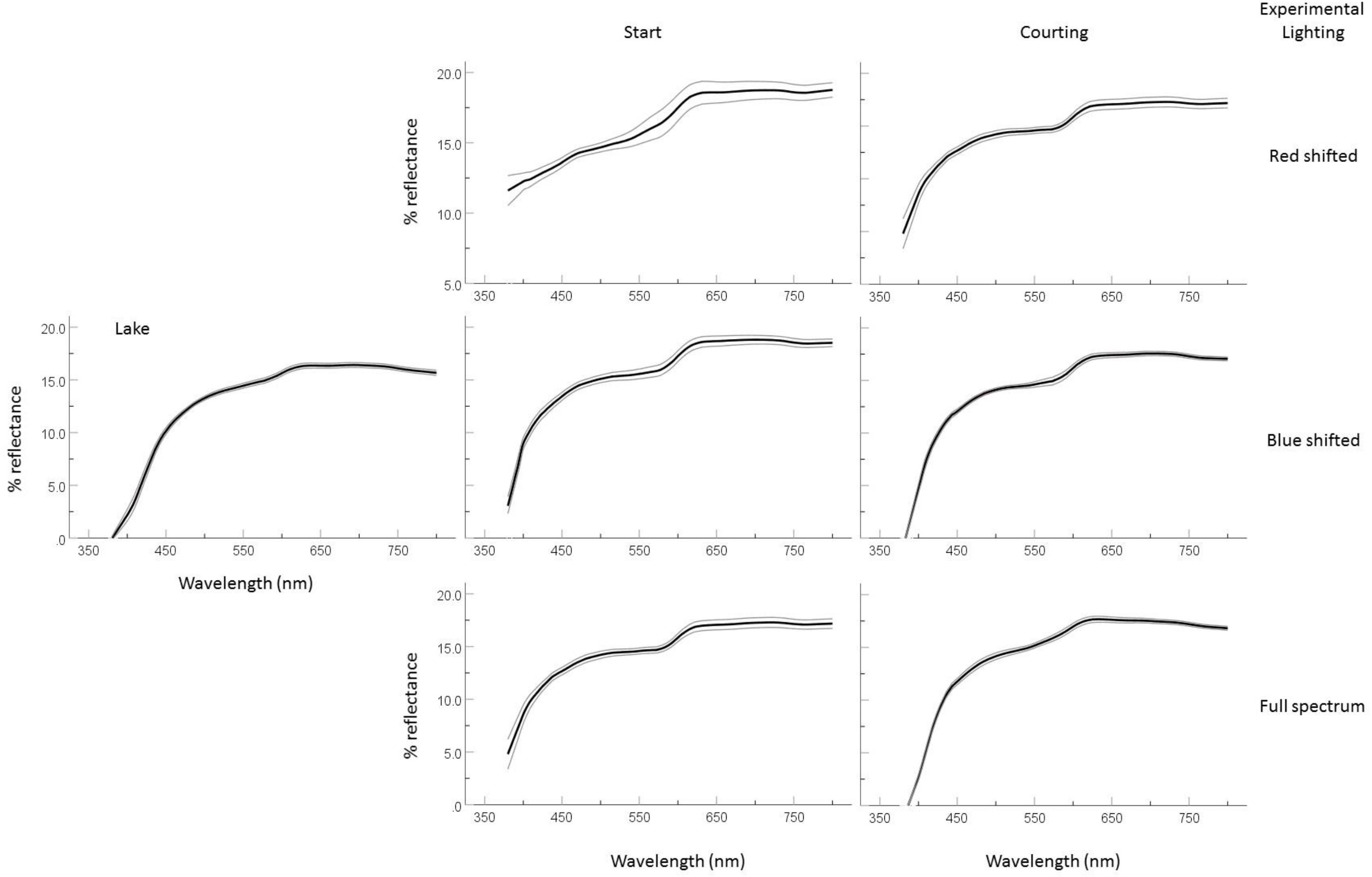
Mean (+/- 95% confidence band) reflectance curves (% reflectance relative to a standard; Ocean Optics WS-1) for males at time of capture and at experimental day 0 and day 13 (courting).

**Figure 3.**
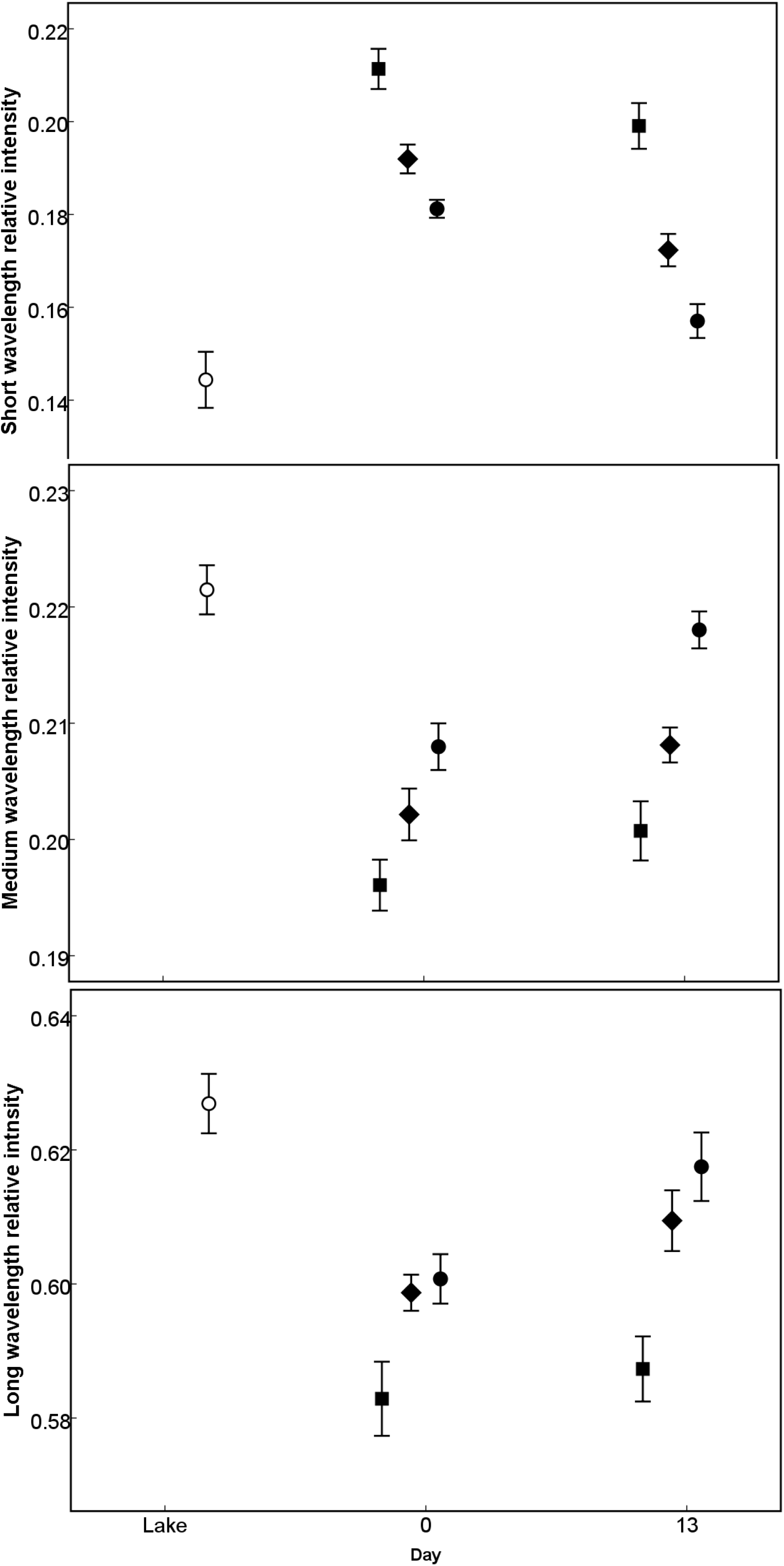
Relative intensity of short, medium and long wavelengths of light reflected from male nuptial signals 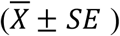 for male threespine stickleback collected from Hotel Lake (Day = “Lake”) and subsequently held in each of the three lighting treatments (square – red shifted, diamond – full spectrum reduced intensity and circles – blue-green shifted) at the start (Day=1) and end (Day = courting) of the experiment.

## Discussion

The present study demonstrates that the male threespine stickleback nuptial signal is plastic, responding to both the change in ambient lighting from the wild to the laboratory, and to different experimental lighting environments. Males across all lighting treatments increased the relative intensity of short-wavelength reflectance and decreased the relative intensity of medium and long-wave length reflectance of their signal following transfer from the wild to the laboratory. Males nesting in the red-shifted treatment had higher relative reflectance in the short wave-length range of the spectrum compared to the males in both the blue-green shifted and full-spectrum treatments, and had lower relative reflectance in the medium and long wavelength regions of the spectrum compared to males nesting in the other two treatments as well. The nature of these changes is similar to those observed by Brock et al. (2017) in their investigation of changes in male stickleback signal with changes in ambient light associated with a signalling male’s nest depth.

Plasticity of the male threespine stickleback nuptial signal has been demonstrated previously in a few other studies. For example, Barber et al. (2000) found that both the red component of the signal and the signal intensity increased following transfer from field lighting to laboratory lighting. Hulslander (2003) found that signals become over-expressed when individuals from some Alaskan populations are moved from the wild into artificial lighting conditions. Finally, rearing (embryo to adult) environment (full-spectrum or red-shifted light) influences the signal characteristics of melanic stickleback from Enos Lake, BC, mosaic stickleback from Paxton Lake, BC and hybrids between these two colour morphs (Lewandowski and Boughman 2008).

Modification of integument reflectance patterns under various environmental, social, and stressful conditions is common among teleost fish (Rush et al. 2003, Amiya et al. 2005, Scott 2011, Costa et al. 2017), and plastic responses to variability in habitat specific lighting conditions has been well-establish (e.g. Donnelly and Whoriskey 1991, Ellis et al. 1997, Kekalainen et al. 2010, Clarke and Schluter 2011, Nyboer et al. 2014). Reflectance patterns can be altered rapidly by changing the distribution of pigments within a chromatophore (physiological colour change) or slowly by changing the density of various types of chromatophores (morphological colour change; Fujii 2000, Sugimoto 2002, Nyboer et al. 2014). I cannot tell which mechanism is responsible for the variability observed in the present study.

Carotenoid pigments make up the striking red nuptial signals of many species (Anderson 1994; Olsen and Owens 1998; Wedekind et al. 1998), and have been implicated in signalling an individual’s quality as a mate (Hill 1992; Lozano 1994; Hill 2000; Alonso-Alverez et al. 2004; McGraw 2005). In fish, red-based signals are commonly achieved by deposition of carotenoid pigments in the phospholipid layer of dermal cells (Svensson and Wong 2011) and therefore are not rapidly mobilized. However, once carotenoid based signals have been established, their appearance can be modified by the dispersion of melanin in the overlying melanophores, which can occur rapidly (Fujii 2000, Sugimoto 2002). Fish producing a red mating signal, such as male threespine stickleback, may respond plastically to variability in ambient light by adjusting carotenoid pigment deposition or by altering the distribution of melanin. Though Laurin and Scott (2009) showed that the male threespine stickleback nuptial signal is relatively stable for up to 20 minutes following transfer to artificial lighting, the present study demonstrates that the threespine stickleback nuptial signal will respond to lighting changes in as little as one day.

Male threespine stickleback nuptial signal expression responds to variability in dietary carotenoid content in the lab (Pike et al. 2007, 2011; but see Scott and Black 2018), and variability among male threespine stickleback in nuptial signal expression has been correlated with variability in fitness; males with redder, more carotenoid rich signals tend to be preferred by females to, and are better parents than males with less red, less carotenoid rich signals (Bakker 1990, Bakker and Mundwiller 1994, Rowland 1994, Pike 2007a, b). All fish used in the present study had access to natural levels of dietary carotenoids prior to collection and were fed a common diet of frozen brine shrimp following transfer to the laboratory. Differences among individuals in stored carotenoids and consequently to variability in nuptial signal reflectance characteristics is unlikely to have influenced the results of the present study because (i) signal reflectance characteristics do not correlate with stored or dietary carotenoid pigments in this population (Black et al. 2014; Scott and Black 2018), and (ii) the randomization process would have eliminated any impact of variability among males due to variability produced *in situ*. Therefore, the signal reflectance patterns observed in this study were a result of alteration of the lighting environment only.

The majority of studies that have examined the stickleback nuptial signal have been conducted under artificial light (Milinski and Bakker 1990, Pike et al 2007a, 2011, Rick et al. 2011, Hiermes et al. 2016). Mate choice trials set in more realistic lighting environments often lead to different results compared to natural lighting (Hughes et al. 2013); indeed, female stickleback from red populations show no colour-based preference for males under natural lighting conditions (Scott 2004). The results of the present investigation and of other stickleback studies demonstrating signal plasticity in response to ambient light variability (eg. Brock et al. 2017; Scott and Black 2018), suggest that standard stickleback mating experiments should be repeated under more realistic lighting conditions.

Natural selection should favour phenotypic plasticity in species that colonize new environments, or that inhabit temporally or geographically variable environments (Foster 1995, Via et al. 1995, Pigliucci 2005, Volis et al. 2015). Threespine stickleback mate in a variety of systems that vary in intensity and composition of various wavelengths of light, including coastal marine, estuarine, and brackish systems as well as oligotrphoic, eutrophic, dystrophic, and lotic freshwater systems. Spectral properties of light in water alters the efficacy of visual signals (Fuller and Noa 2012, Hueschele et al. 2012, Giery and Layman 2017), and variability among systems in these properties will favour signals that can respond plastically and maintain their effectiveness across various habitats (Endler 2000, Chapman 2009, van der Sluijs et al. 2011, Barquerro et al. 2015). The threespine stickleback mating signal responds plastically to various lighting environments as is expected given the species’ ability to colonize new habitat, its ability to thrive in temporally and spatially variable habitats, and the variable nature of the spectral environments it colonizes. Phenotypic plasticity can facilitate rapid adaptation to novel environments and may enable rapid speciation since it provides colonizers the ability to establish populations in new and diverse environments (the initial stage is dependent upon plasticity) followed by isolation and genetic divergence/ adaptation (Wund et al. 2008; Pfennig et al. 2010; Young 2013; Schneider and Meyer 2017). Plasticity in traits related to mate recognition may be especially good at producing reproductive isolation (Thibert-Plante and Hendry 2011) since they can lead to rapid establishment of signal based assortative mating (Scott 2004).

Here, I demonstrated plasticity of the male stickleback nuptial signal in response to variation in ambient light. The prominent role that investigations of this signal have played in developing our understanding of behavioural ecology, especially with regard to sexual selection and honest signalling in this species and in general brings forward the question of how relevant the laboratory investigations have actually been. Have we built a set of empirical generalities based on an artifact of the laboratory setting? Clearly, studies of stickleback signalling dynamics need to be repeated under realistic ambient lighting conditions, at the very least to validate what we already think we know.

## Acknowledgements

Valerie Mucciarrelli and Diana Lawry assisted with collection and maintenance of stickleback as well as making irradiance and reflectance measurements. Michael Jackson, Ruby Lake Lagoon Society, granted access to the Iris Griffith Field Studies & Interpretive Centre in Medeira Park, BC, Canada for conducting the lighting experiment. Funding for the research was provided by an NSERC Discovery Grant and the NSERC USRA program. This research was reviewed and approved by the Animal Use Sub-committee of the Animal Care Committee at Western University, London, Ontario, Canada.

## Notes

### Competing Interest Statement

The authors have declared no competing interest.

